# High resolution genome-wide occupancy in *Candida spp*. using ChEC-seq

**DOI:** 10.1101/2020.06.19.160531

**Authors:** Faiza Tebbji, Inès Khemiri, Adnane Sellam

## Abstract

To persist in their hostile and dynamic human host environments, fungal pathogens has to sense and adapt by modulating their gene expression to fulfil their cellular needs. Understanding transcriptional regulation on a global scale would uncover cellular processes linked to persistence and virulence mechanisms that could be targeted for antifungal therapeutics. Infections associated with the yeast *Candida albicans*, a highly prevalent fungal pathogen, and the multi-resistant related species *C. auris* are becoming a serious public health threat. To define the set of a gene regulated by a transcriptional regulator in *C. albicans*, Chromatin Immuno-Precipitation (ChIP) based techniques including ChIP-chip or ChIP-seq has been widely used. Here, we describe a new set of PCR-based MNase-tagging plasmids for *C. albicans* and other *Candida spp*. to determine genome-wide location of any transcriptional regulator of interest using Chromatin endogenous cleavage (ChEC) coupled to high-throughput sequencing (ChEC-seq). The ChEC procedure does not require protein-DNA crosslinking or sonication avoiding thus artefacts related to epitope masking or the hyper-ChIPable euchromatic phenomenon. In a proof-of-concept application of ChEC-seq, we provided a high-resolution binding map of the SWI/SNF chromatin remodeling complex, a master regulator of fungal fitness in *C. albicans* in addition to the transcription factor NsiI that is an ortholog of the DNA-binding protein Reb1 for which genome-wide occupancy were previously established in *Saccharomyces cerevisiae*. The ChEC-seq procedure described here will allow a high-resolution genomic location definition which will enable a better understanding of transcriptional regulatory circuits that govern fungal fitness and drug resistance in these medically important fungi.

**Importance:** Systemic fungal infections caused by *Candida albicans* and the ‘superbug’ *C. auris* are becoming a serious public health threat. The ability of these yeasts to cause disease is linked to their faculty to modulate the expression of genes that mediate their escape from the immune surveillance and their persistence in the different unfavourable niches within the host. Comprehensive knowledge on gene expression control of fungal fitness is consequently an interesting framework for the identification of essential infection processes that could be hindered by chemicals as potential therapeutics. Here, we expanded the use of ChEC-seq, a technique that was initially developed in the yeast model *Saccharomyces cerevisiae* to identify genes that are modulated by a transcriptional regulator, to the pathogenic yeasts *C. albicans* and *C. auris*. This robust technique will allow a better characterization of key gene expression regulators and their contribution to virulence and antifungal resistance in these pathogenic yeasts.

## Introduction

*Candida* species, in particular *Candida albicans*, are major components of the disease burden caused by fungi and are frequent cause of life-threatening invasive infections especially in immunocompromised patients. The emergent *C. auris* was the first fungal pathogen considered as an urgent public health threat due to its multidrug resistance, high transmissibility among patients in health-care facilities and elevated crude mortality (1). Other *Candida* species such as *C. parapsilosis, C. tropicalis, C. guilliermondii* and the azole-resistant yeasts *C. glabrata* and *C. krusei* are also frequent cause of Candidiasis and vulvovaginal infections (2–4). Current anti-*Candida* therapeutics suffer from diverse limitations including toxicity, resistance and interactions with other commonly prescribed drugs. This has led to increasing interest in studying mechanisms underlying resistance and virulence of *Candida* species with the ultimate goal to identify potential drug targets for novel antifungal therapeutic intervention. However, the diploid nature and the absence of a complete sexual cycle in most of *Candida* species limit the use of classical genetic approaches to dissect mechanisms controlling fungal fitness and antifungal resistance. Alternatively, applying genome-wide transcriptional methods such as those determining gene expression changes (DNA microarrays and RNA-seq) or genomic occupancy (ChIP-chip and ChIP-seq) in *Candida* species had significantly contributed to uncovering different facets of fungal biology that are critical for both opportunistic and commensal lifestyles, in addition to antifungal tolerance and resistance (5–17). These approaches had also helped to uncover a surprising extent of evolutionary plasticity of transcriptional regulatory circuits in these species as compared to the model yeast *S. cerevisiae* (18, 19).

While ChIP-chip and ChIP-seq have been traditionally used to unbiasedly map the binding of a transcriptional regulator (TR), this tool has some limitations that are attributed mainly to TR-DNA crosslinking and DNA shearing by sonication (20). Formaldehyde is commonly used for Protein-DNA crosslinking, however, this chemical preferentially generates protein-protein crosslinks which can cause epitope masking and consequently alters the efficiency of the immunoprecipitation procedure and led to increased signal background noise. Furthermore, DNA fragmentation by sonication can disrupt weak or transient TR-DNA or TR-histone interactions and generate DNA fragments with heterogenous sizes and thus, impede the refinement of binding site identification (21).

To circumvent these limitations, crosslinking- and sonication-free alternative methods has been developed recently (20, 22–25). In one such method termed chromatin endogenous cleavage (ChEC) (26), the TR of interest is fused to the micrococcal nuclease (MNase) in order to fragment unprotected neighboring chromatin upon MNase-activation by calcium (26). ChEC coupled to high-throughput sequencing (ChEC-seq) was efficiently used to map bindings of the generalist transcription factors Reb1, Abf1 and Rap1 in the budding yeast and has provided a high-resolution occupancy with more binding events as compared to ChIP-based tools (20). Additionally, temporal analysis of ChEC-seq data uncovered that TR can have two distinct binding behaviors; a fast binding uncovered by rapid MNase cleavage at locus with robust *bona fide* TR-binding motif and a second slow cleavage with low-scoring motifs that are preferentially sampled by a given TR. ChEC-seq has been successfully used to define genomic occupancy of the chromatin remodeler RSC complex (Rsc8 subunit) as ChIP procedure was less efficient (27, 28). Several recent investigation took advantage from ChEC-seq to study the role of different core components of the general transcriptional machinery such as mediators, SAGA, histone acetyltransferases and chromatin “pushers”, on global gene expression control and promoter nucleosome architecture in eukaryotes (29–32).

In this work, we describe a new set of PCR-based MNase-tagging plasmids for *C. albicans* and other *Candida* species to determine genome-wide location of any TR of interest by ChEC-seq. In a proof-of-concept application of ChEC-seq in *C. albicans*, we have selected NsiI that is an ortholog of the DNA-binding protein Reb1 for which genome-wide occupancy were previously established by ChEC-seq in *S. cerevisiae* (20). As our previous effort on mapping occupancy of the *C. albicans* chromatin remodeling complex SWI/SNF by ChIP-tiling arrays had led to a substantial background noise (7), we have used the ChEC-seq assay to obtain a high-resolution binding map of this master regulator of fungal fitness (6). The ChEC-seq procedure described here will allow a high-resolution genomic location definition which will enable a better understanding of transcriptional regulatory circuits that govern fungal fitness and drug resistance in these medically important fungi.

## Methods

### Strains and media

*C. albicans* was routinely maintained at 30°C on YPD (1 % yeast extract, 2 % peptone, 2 % dextrose, with 50 mg/ml uridine). The *C. albicans* WT strain SN148 (*his1/his1, leu2/leu2, arg4/arg4, ura3*/*ura3*::imm434, *IRO1/iro1*::imm434) (33) used in this study derives from the SC5314 clinical strain. For *C. auris*, the clinical B8441 strain (34) was used for *SAT1*-MNase-tagging. For spot dilution assays, overnight cultures of both *C. albicans* and *C. auris* were diluted to an OD_600_ of 1 and fivefold serial dilutions were prepared in distilled water. A total of 4 μl of each dilution were spotted on YPD-agar plates for 1 days at different temperatures (30 □C, 37°C and 40°C) and imaged using the SP-imager system.

### Construction of the pFA-MNase plasmids and the “MNase free” control strain

The pFA-MNanse-CaURA3, pFA-MNase-CaHIS1 and pFA-MNase-CaARG4 plasmids were constructed as following: DNA of the 3x FLAG epitope-MNase module was synthesized by Biobasic, with codon optimized for *C. albicans* (a total of eleven CTG codons of the MNase were changed to TTA or TTG). The *PacI-AscI* 3x FLAG-MNase fragment was cloned in the *PacI-AscI* digested pFA-TAP-CaURA3, pFA-TAP-CaHIS1 and pFA-TAP-CaARG4 (35). For pFA-MNase-SAT1, the pFA-MNase-Ca*URA3* was double digested with *AscI* and *Sac1* restriction enzymes to remove the *URA3* auxotrophy marker. *SAT1* marker was amplified from pFA-SAT1 (36) with primers (**Table S1**) containing restriction sites *AscI-Sac1* and cloned into the *AscI-Sac1* digested pFA-MNase. The resulting pFA-MNase-SAT1 was sequenced to confirm the integrity of the Sat1 dominant marker.

The MNase-free control strain was constructed as following: The *C. albicans* codon-optimized DNA of the 3xFLAG-Mnase-nuclear localization signal (SV40) construct delimited by *NheI* and *MluI* restriction sites was synthesized. The *NheI-MluI* digested 3xFLAG-Mnase-SV40 fragment was then cloned into CIp-p*ACT1*-CYC vector (37) to ensure constitutive expression of MNase in *C. albicans*. The CIp-p*ACT1*-3xFLAG-Mnase-SV40-CYC plasmid was linearized by *StuI* restriction enzyme and integrated at the *RPS1* locus of the SN148 WT strain.

For the *C. auris* MNase control strain, the *URA3* auxotrophy marker of the *CIp-pACT1*- 3xFLAG-Mnase-SV40-CYC was replaced by SAT1 as follow: *SAT1* marker was amplified from pFA-SAT1 with primers containing restriction sites *NotI-NheI* (**Table S1**) and cloned into the *NotI-NheI* digested CIp-p*ACT1*-3xFLAG-Mnase-SV40-CYC. To allow the integration of the CIp-p*ACT1*-3xFLAG-Mnase-SV40-CYC in the *C. auris* genome, the *C. albicans RPS1* integrative locus was replaced by a short 900-bp *C. auris* intergenic region *CauNI* (*<u>C. aur</u>is* Neutral Intergenic; PEKT02000001: 871,442-872,342) as following: *CauNI* region was amplified from the *C. auris* B8441 genomic DNA with primers containing restriction sites *NotI*-*NheI* (**Table S1)** and cloned into the *NotI*-*NheI* digested CIp-p*ACT1*-3xFLAG-Mnase-SV40-CYC-*SAT1* plasmid. The resulting plasmid was linearized by *StuI* restriction enzyme and integrated at the *CauNI* locus of the *C. auris* B8441 strain. The correct integration of the CIp-p*ACT1*-3xFLAG-Mnase-SV40-CYC-*SAT1* cassette was verified by PCR. Integration of any exogenous DNA or the CIp-p*ACT1*-3xFLAG-Mnase-SV40-CYC-*SAT1* at the *CauNI* has no impact on the *in vitro* growth of *C. auris*.

### PCR-based tagging of endogenous loci in *C. albicans* and *C. auris*

*SNF2* (C2_02100W_A) and *NSI1* (C6_03550C_A) were Mnase-tagged *in vivo* with the Mnase cassette PCR products following the protocol described by Lavoie *et al*. (35). The MNase cassettes were amplified using 120-bp primer pair with 20 bp of vector sequences (Forward: GGTCGACGGATCCCCGGGTT and Reverse: TCGATGAATTCGAGCTCGTT) and 100 bp from *SNF2* (C2_02100W_A) and *NSI1* (C6_03550C_A) (**Table S1**). PCR reactions were performed in 50 μl volumes with 1 ng pFA-MNase plasmid and the Q5 high fidelity polymerase (New England Biolabs). PCR thermocycling was executed as following: initial denaturing, 98°C for 3 min; 35 cycles using 98°C for 10 sec, 56°C for 30 sec and 72°C for 3 min. PCR products were used directly to transform the WT strain SN148 using lithium acetate transformation protocol (38). Transformants were selected on selective plates and positive colonies were analyzed by PCR to confirm the correct integration of the MNase-tag. For *C. auris*, *CauSNF2* (B9J08_001192) and *CauNSI1* (B9J08_003000) were both Mnase-tagged *in vivo* with the Mnase-*SAT1* cassette as described for *C. albicans* with the exception that the 20 bp of the reverse vector sequence was: TCTGATATCATCGATGAATTCGAG.

### ChEC-seq procedure

For each ChEC experiment, saturated overnight cultures of *C. albicans* Mnase tagged and Mnase free strains were diluted to a starting OD_600_ of 0.1 in 50 ml YPD medium. Cells were grown at 30°C to an OD_600_ of 0.7–0.8. Cells were pelleted at 3,000 g during 5 min and washed three times with 1 ml Buffer A (15 mM Tris pH 7.5, 80 mM KCl, 0.1 mM EGTA, 0.2 mM spermine, 0.5 mM spermidine, one tablet Roche cOmplete EDTA-free mini protease inhibitors, 1 mM PMSF). Cells were then resuspended in 800 μl Buffer A containing 0.1% digitonin (Sigma) and permeabilized for 10 min at 30 °C under shaking. MNase digestions were performed by adding CaCl2 at final concentration of 5 mM and incubated at the indicated time at 30 °C. At each time point, a total of 200 μl aliquots of the ChEC digestions were transferred to a tube containing 50 μl of 250 mM EGTA to quench MNase digestions. For each factor analyzed, the timepoint “0” corresponds to a condition where MNase were not activated by CaCl2. Nucleic acids were extracted using MasterPure yeast DNA purification kit (Epicentre, MPY80200) according the manufacturer’ instructions, and resuspended in 50 μl 0.1 X Tris-HCl buffer, pH 8.0. RNAs were digested with 10 μg RNase A at 37 °C for 20 min. To assess MNase activity, 5 μl of digested DNA of each ChEC time-point (time after CaCl_2_ addition) was loaded on 1.5% agarose gel. ChEC DNA was subjected to size selection using the Pippin Prep (SageScience) size-selection system with 2% agarose gel cassette allowing the removal of multi-kilobase genomic DNA fragments and the enrichment of 100-400 bp DNA fragments.

### Library preparation, NGS sequencing and peak calling

The NEBNext Ultra II DNA Library Prep Kit for Illumina was used to construct the ChEC-seq library following the manufacturer’s instruction. The quality, quantity and the size distribution of the libraries were determined using an Agilent Bioanalyzer. A 50-bp paired-end sequencing of DNAs were performed using an Illumina HiSeq 4000 sequencing system. Sequence were trimmed to remove adapters using TRIMMOMATIC (39). Read ends were considered to be MNase cuts and were mapped to the *C. albicans* genome (*Candida_albicans_SC5314* assembly 22) (40) using Bowtie2 (41). ChEC alignment and track visualization using bedgraph files were performed as previously described (20, 42). Peaks were determined from the normalized ChEC ratio using MACS2 algorithm (43) with a window size of 60 bp. Cis-regulatory motif enrichment was assessed in the top high scoring 1000 peaks for both Nsi1 and Snf2 using MEME-ChIP software (44).

### Data availability

The sequences of plasmids pFA-MNase-CaHIS1, pFA-MNase-CaARG4, pFA-MNanse-CaURA3 and pFA-MNase-SAT1 have been submitted to GenBank and have been assigned the following accession numbers: MT181237, MT181238, MT181239 and MT223485. All ChEC-seq data generated in this study were submitted to GEO database under the accession number GSE150063.

## Results and discussion

### Plasmid toolbox for MNase tagging in *C. albicans* and *non-albicans Candida* species

We have previously constructed a series of pFA plasmids for C-terminal HA-, TAP- and MYC-tagging in *C. albicans* with the *URA3, HIS1* and *ARG4* autotrophy markers (35). Here, we have used these plasmids as a starting point to build a new pFAs plasmids that allow a C-terminal tagging of any protein of interest at its native chromosomal location with the MNase. We synthesized a DNA construct encoding a 3x FLAG epitope and MNase that have been codon-optimized for *C. albicans*. This construct was used to replace the DNA sequence of the TAP-tag in the pFA-TAP-CaURA3, pFA-TAP-CaHIS1 and pFA-TAP-CaARG4 to generate the pFA-MNanse-CaURA3, pFA-MNase-CaHIS1 and pFA-MNase-CaARG4, respectively. These plasmids allow the use of a single 120-bp primer pair (20 bp of vector sequences and 100 bp from the gene to be tagged) for PCR-based tagging of endogenous loci in *C. albicans* (**Figure 1A**). These primers are compatible with the pFA-TAP/HA/MYC (35) and the pFA-XFP tagging systems (36, 45).

**Figure 1.**
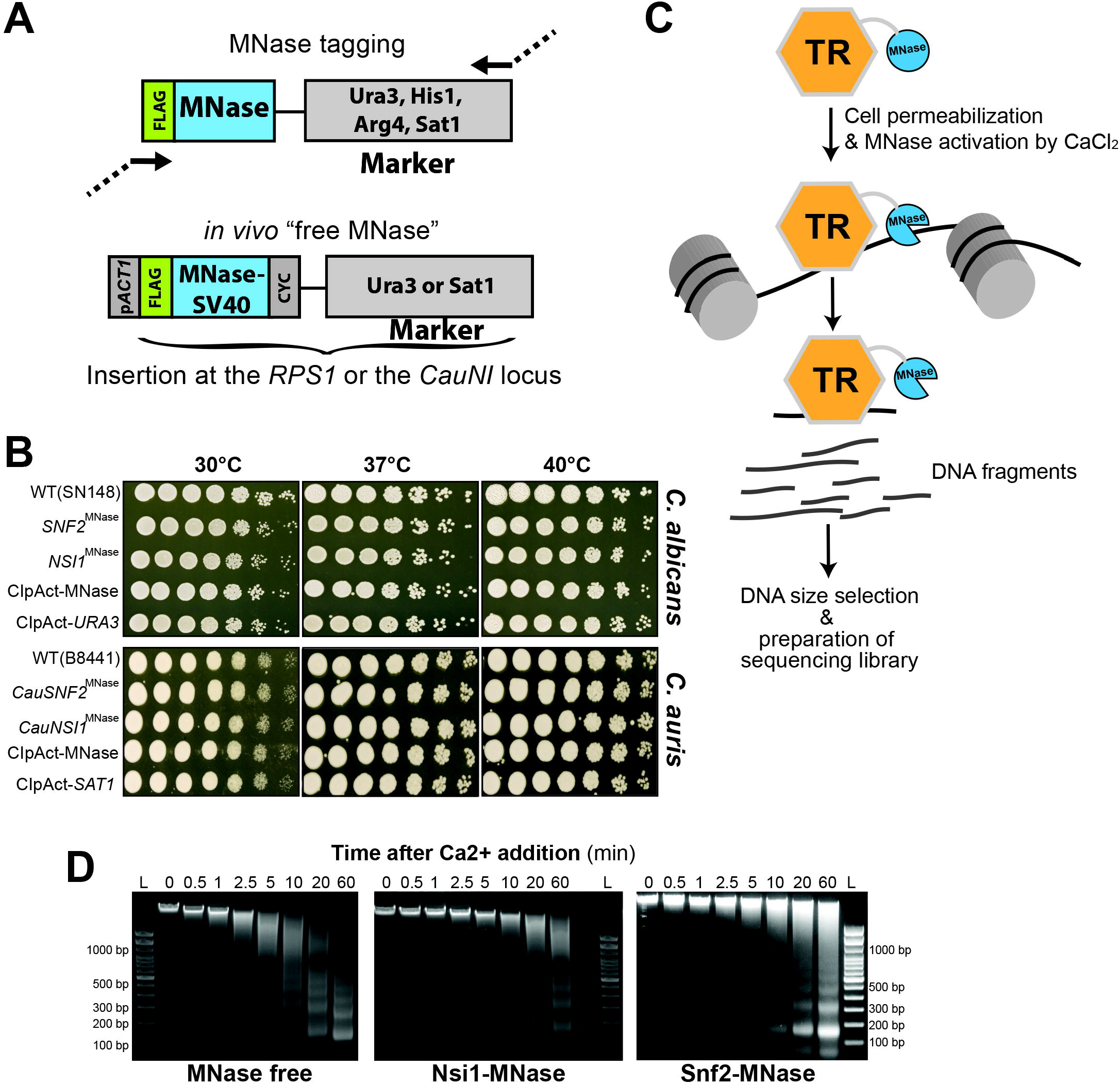
ChEC-seq method in *C. albicans* and *Candida spp*. **(A)** Plasmid constructs for *in vivo* TR-MNase tagging and the construction of the “free MNase” control strains in *C. albicans* and *C. auris*. **(B)** Phenotypic characterization of strains bearing the MNase-tagged Nsi1 and Snf2 TRs, and the “free MNase” control constructs in *C. albicans* and *C. auris*. For both species, WT, Snf2^MNase^, Nsi1^MNase^, the free MNase and the control (empty vector) strains were serially diluted, spotted on YPD and incubated for one day at different temperatures. **(C)** Schematic representation of the experimental setup of the ChEC-seq methodology. *Candida spp*. cells where a TR of interest is fused to MNase are permeabilized with digitonin prior to MNase activation with calcium. This will lead to the fragmentation of unprotected neighboring chromatin. The resulting fragmented DNA is purified and subjected to size selection prior to high-throughput sequencing. **(D)** Evaluation of genomic DNA fragmentation by agarose gel electrophoresis at 0, 0.5, 1, 2.5, 5, 10, 20 and 60 minutes of calcium exposure in the Snf2^MNase^, Nsi1^MNase^ and the free MNase strains in *C. albicans*. L: DNA ladder.

We have also constructed the pFA-Mnase-SAT1 plasmid with the dominant selectable marker *SAT1* that confers resistance to the antibiotic nourseothricin for MNase-tagging in clinical strains of *C. albicans* and non-*albicans Candida* species such as the superbug *C. auris*.

### ChEC-seq experimental procedure

ChEC-seq has been initially used in *S. cerevisiae* to map the genomic occupancy of canonical general regulatory factors such as the RNA polymerase I Enhancer Binding protein ScReb1 (20). Here, we have selected the Nsi1 (C6_03550C_A), that is the ScReb1 ortholog in *C. albicans*, to perform ChEC-seq. Additionally, we were also interested in the catalytic subunit of the SWI/SNF complex, Snf2, to explore the potential of ChEC-seq in mapping genomic occupancy of chromatin remodeling complexes. We have previously mapped the genomic location of the *C. albicans* Snf6 that is a fungus specific SWI/SNF subunit using ChIP coupled to high-density tiling arrays (7). Thus, the SWI/SNF genome-wide binding data generated by ChEC-seq can be compared to those of ChIP-chip to assess the sensitivity of each technique. We also generated an MNase control strain (“free MNase”) with a 3xFLAG-tagged MNase module fused to an SV40 nuclear localization signal under the control of *ACT1* and integrated at the *RPS1* locus. Constitutive expression of MNase (Free MNase) or MNase-tagging of Nsi1 or Snf2 had no perceptible effect on the growth of *C. albicans* at different temperatures (**Figure 1B**).

Both *C. auris* Snf2 and Nsi1 orthologs were also MNase-tagged using PCR cassettes generated from the pFA-Mnase-SAT1 plasmid. For the MNase control strain, 3xFLAG-tagged MNase module was inserted into the neutral intergenic locus *CauNI* where integration has no effect on the *in vitro* fitness of *C. auris* (**Figure 1B**). As for *C. albicans*, MNase-tagging of CauNsi1 and CauSnf2 does not affect the growth of *C. auris* (**Figure 1B**).

To provide a proof of principle for using ChEC in *Candida spp*., we focused our effort on *C. albicans*. The *S. cerevisiae* ChEC procedure described by Zentner *et al*. (20) (**Figure 1C**) was followed with some modifications. *C. albicans* cells were permeabilized with digitonin for 10 min prior to MNase activation with 5mM CaCl_2_. We presumed that the treatment of permeabilized cells with calcium would engender both specific and non-specific cleavages. We therefore made a size selection of the ChEC DNA before preparing the sequencing library to enrich small fragments less than 400 bp. Prior to size selection and for each transcriptional regulator and the free MNase strain, we analyzed the kinetic of DNA digestions by agarose gel electrophoresis. Analysis of minute-scale timepoints revealed notable smearing of genomic DNA of all TF-MNase fusions by the 5 min time point. This pattern increased over time until 60 min. In contrast, digestion in the free MNase strain yielded earlier a smearing by 30 s (**Figure 1D)**. The 5, 20 and 60 min digestion times were selected for both Snf2 and Nsi1 ChEC-seq experiments. Size selection was performed using the Pippin Prep size-selection system with 2% agarose gel cassette. The goal of this stage is to remove multi-kilobase fragments of genomic DNA and enriched a small fragment. The 2% Agarose gel cassettes allows enrichment of DNA fragments below 100-400 bp. Alternatively, size selection could be performed using the paramagnetic beads for size selection and buffer exchange steps such as the AMPure XP cleanup kit (Beckman coulter) (20, 46).

### Genome-wide binding of Nsi1 and Snf2 by ChEC-seq

To assess the ChEC-seq performance in *C. albicans*, we have chosen to map the genomic occupancy of NsiI that is an ortholog of the DNA-binding protein Reb1 for which the genomewide occupancy were previously established by ChEC-seq in *S. cerevisiae* (20). We detected 2548, 4771 and 4523 NsiI peaks upon 5, 20 and 60 min MNase activation, respectively (**Table S2 and Figure 2A**). *De novo* motif analysis of intergenic bound Nsi1 regions showed a significant enrichment of the *bona fide* Reb1/Nsi1 site (TTACCCGG) at 5 min while a non-specific long AC/TG-rich sequence was the most enriched at 20 and 60 min (**Figure 2B**). This suggest that the 5 min MNase cleavage mapped the Nsi1 fast class binding events while 20 and 60 min correspond to the slow class binding that lack robust consensus motif. Thus, as for *S. cerevisiae*, our ChEC-seq data recapitulated the time-dependent binding behavior of transcriptional regulators and can be used to map early high-affinity interactions with consensus motifs and sequence that are preferentially sampled by a given protein (20). Our genome-wide occupancy data recapitulated the overall functions of either Nsi1 or Reb1 in *S. cerevisiae* as reflected by Nsi1 binding to the promoter of ribosome biogenesis and rRNA genes (**Figure 2C-F**) (47–49).

**Figure 2.**
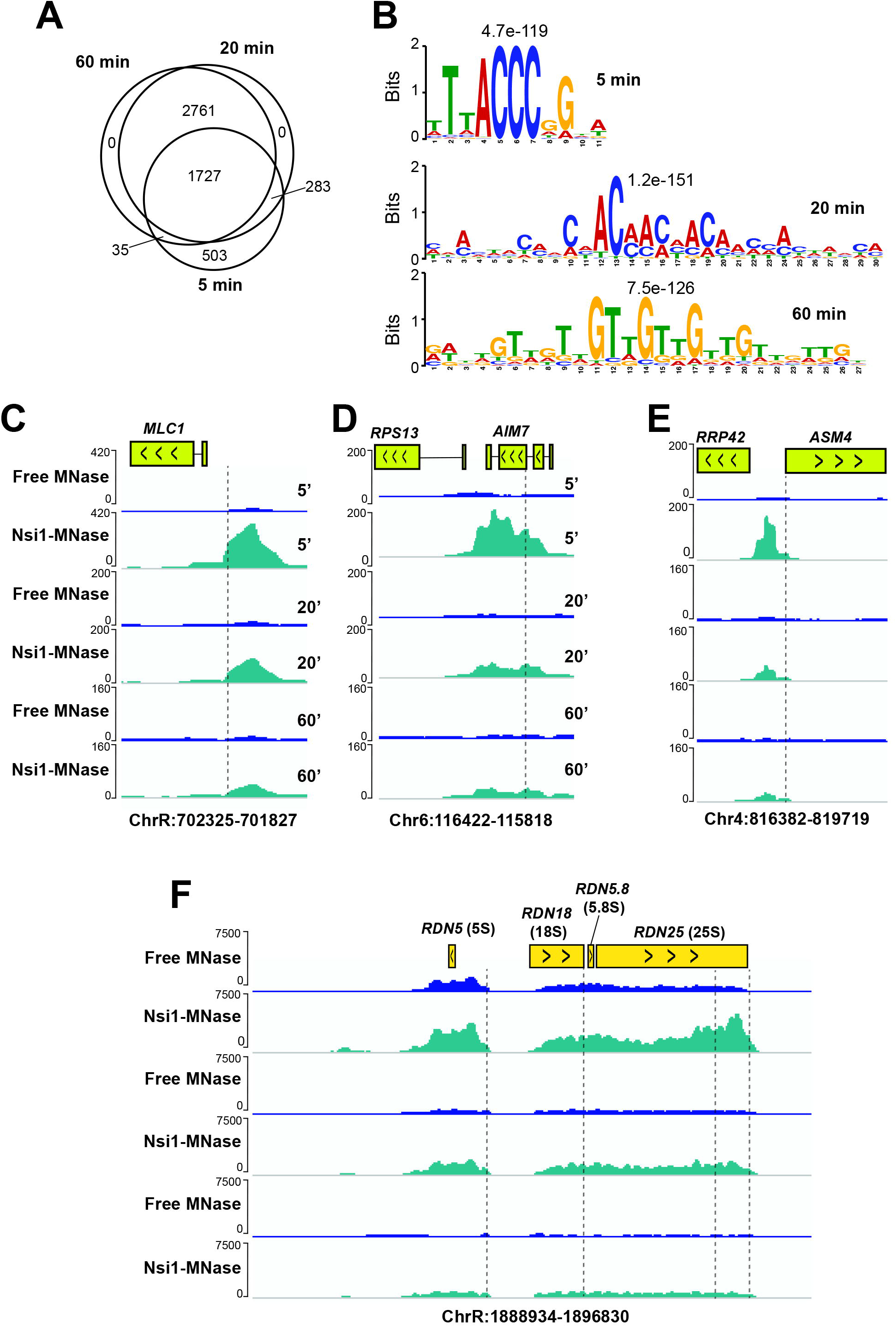
Genome-wide occupancy of the transcription factor Nsi1 with ChEC-seq. **(A)** Temporal analysis of Nsi11 binding events. Venn diagram showing the overlap of Nsi1 binding events at three distinct MNase activation times (5, 20 and 60 min). **(B)** Motif scores for Nsi1 bound promoters at 5, 20 and 60 min after MNase activation. The motif logos were generated using MEME-ChIP software on the 1000 high scoring peaks. **(C-F)** Snapshot of genomic regions showing the ChEC-seq signal for Nsi1^MNase^ and the free MNase strains at 5, 20 and 60 min after MNase activation. The position of Nsi1 motifs are indicated by the dashed lines. Nsi1 occupies the promoter of *MLC1* (**C**), *AIM7* (**D**) and *ASM4* (**E**), in addition to many sites within the rDNA locus **(F)**.

ChEC-seq of Snf2 identified 4145, 6446 and 6215 peaks at 5, 20 and 60 min MNase cleavage, respectively, which is 10-fold higher than the number of peaks detected under similar growth conditions by ChIP-tiling array of the SWI/SNF subunit, Snf6 (7) (**Figure 3A-B and Table S3**). As for Nsi1, the 20 and 60 min ChEC-seq data were similar and might capture the slow sites. The Snf2 fast bound promoters were enriched mainly in carbohydrate metabolism mirroring the previously characterized role of the SWI/SNF complex in *C. albicans* (**Figure 3C**) (6, 7). Snf2 occupied promoters of hexose transport and carbon utilization genes (galactolysis) that were shown to be modulated by the SWI/SNF subunit Snf5 (6) (**Figure 3D**).

**Figure 3.**
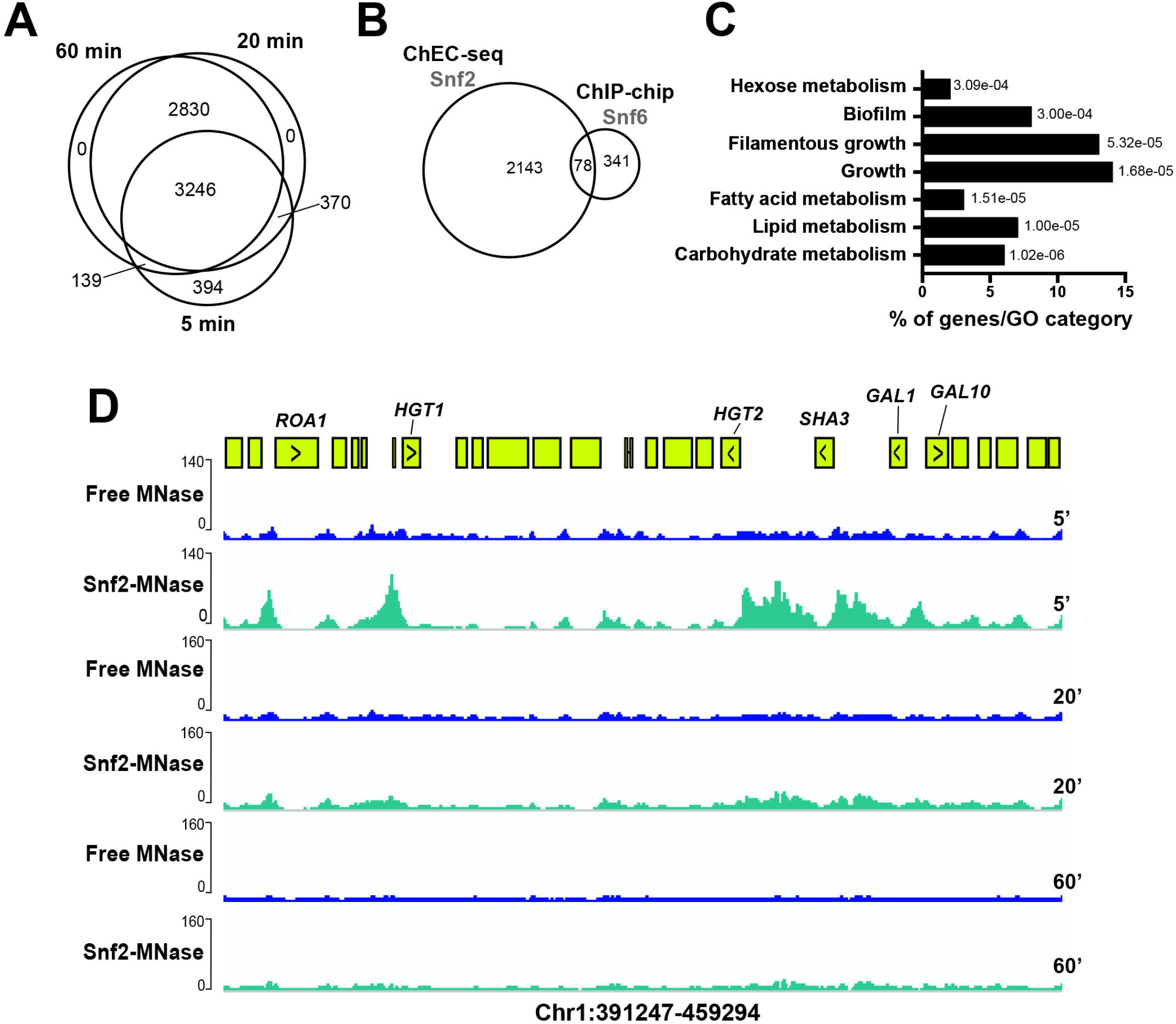
Genome-wide occupancy of the chromatin remodeler Snf2 with ChEC-seq. **(A)** Venn diagram showing the overlap of Snf2 promoter bindings at 5, 20 and 60 min after MNase activation. **(B)** Comparison of SWI/SNF genomic occupancies by ChEC-seq and ChIPchip method. Venn diagram of overlap between Snf2 ChEC-seq sites and Snf6 ChIP-chip using high-density tiling arrays. **(C)** Gene ontology of biological process associated with Snf2-bound promoters at 5 min after MNase activation. The *P* values were calculated using hypergeometric distribution as described in the GO Term Finder Tool website (http://candidagenome.org/cgi-bin/GO/goTermFinder). **(D)** Snapshot of genomic regions showing the ChEC-seq signal for Snf2^MNase^ and the free MNase strains at 5, 20 and 60 min after MNase activation. Genome browser view of Snf2 ChEC-seq displaying promoter occupancies of carbohydrate metabolism genes including galactolysis (*GAL1, GaL10*) and hexose transport (*SHA3, HGT2, HGT1*).

## Conclusion

We have constructed a new set of PCR-based MNase-tagging plasmids to map genomic occupancy of different transcriptional regulators in the human pathogenic yeast *C. albicans* and other *non-albicans Candida* species. Compared the other ChIP-based techniques, the ChEC procedure relies on total DNA extraction instead of chromatin solubilization and does not require protein-DNA crosslinking or sonication avoiding thus artefacts related to epitope masking or the hyper-ChIPable euchromatic phenomenon (50, 51). So far, ChEC has been exclusively used in the model yeast *S. cerevisiae* to map chromatin occupancy of general transcriptional regulators (42), chromatin remodelers (27, 30, 52) and histone modifiers (31, 32) in addition to transcription factors (53, 54). As many transcriptional regulators and chromatin remodelers are key virulence and drug resistance factors in *C. albicans* and other fungi (6, 13, 17, 55–57), the ChEC-seq represents thus an attracting tool to unbiasedly decipher transcriptional regulatory networks of fungal fitness.

## Acknowledgment

We thank Benjamin Albert and David Shore (University of Geneva) for sharing reagents and their technical guidance regarding the ChEC protocol. This work was supported by funds from the Canadian Institutes for Health Research project grant (CIHR, grant IC118460) and the startup funds from the Montréal Heart Institute (MHI) to AS. AS is a recipient of the Fonds de Recherche du Québec-Santé (FRQS) J2 salary award.

## Supplementary data

**Table S1.** List of primers used in this study.

**Table S2.** List of Nsi1 binding peaks

**Table S3.** List of Snf2 binding peaks

## Notes

### Competing Interest Statement

The authors have declared no competing interest.

